# Integrating sequence composition information into microbial diversity analyses with k-mer frequency counting

**DOI:** 10.1101/2024.08.13.607770

**Authors:** Nicholas A. Bokulich

## Abstract

K-mer frequency information in biological sequences is used for a wide range of applications, including taxonomy classification, sequence similarity estimation, and supervised learning. However, in spite of its widespread utility, k-mer counting has been largely neglected for diversity estimation. This work examines the application of k-mer counting for alpha and beta diversity as well as supervised classification from microbiome marker-gene sequencing datasets (16S rRNA gene and full-length fungal ITS sequences). Results demonstrate a close correspondence with phylogenetically aware diversity metrics, and advantages for using k-mer-based metrics for measuring microbial biodiversity in microbiome sequencing surveys. K-mer counting appears to be a suitable and efficient strategy for feature processing prior to diversity estimation as well as supervised learning in microbiome surveys. This allows incorporation of subsequence-level information into diversity estimation without the computational cost of pairwise sequence alignment. K-mer counting is proposed as a complementary approach for feature processing prior to diversity estimation and supervised learning analyses, enabling large-scale reference-free profiling of microbiomes in biogeography, ecology, and biomedical data. A method for k-mer counting from marker-gene sequence data is implemented in the QIIME 2 plugin q2-kmerizer (https://github.com/bokulich-lab/q2-kmerizer).

**Importance:** K-mers are all of the subsequences of length *k* that comprise a sequence. Comparing the frequency of k-mers in DNA sequences yields valuable information about the composition of these sequences and their similarity. This work demonstrates that k-mer frequencies from marker-gene sequence surveys can be used to inform diversity estimates and machine learning predictions that incorporate sequence composition information. Alpha and beta diversity estimates based on k-mer frequencies closely correspond to phylogenetically aware diversity metrics, suggesting that k-mer-based diversity estimates are useful proxy measurements especially when reliable phylogenies are not available, as is often the case for some DNA sequence targets such as for internal transcribed spacer sequences.

## Introduction

Diversity metrics are frequently used in microbiome DNA sequencing surveys to quantify the microbial diversity present in individual samples (alpha diversity) as well as the similarity between samples based on the diversity present (beta diversity), for example in studies of microbial ecology, biogeography, as well as in biomedical research (1, 2). Several diversity metrics have been developed that incorporate phylogenetic information into calculations as a means of weighting different features based on their genetic similarity (3, 4). These are most frequently applied in high-throughput marker-gene sequencing surveys of 16S rRNA gene sequences. However, application of these methods can be hampered by some methodological limitations, e.g., the limited phylogenetic accuracy of very short sequence reads (e.g., from Illumina devices) that are frequently used for DNA sequencing but can lead to erroneous phylogenies (5).

Phylogeny-aware diversity metrics can also be challenging to use with some target domains, for example the internal transcribed spacer (ITS) domain, which is a non-coding region located between the small and large subunit rRNA genes, and is most frequently used for molecular classification of fungi, being considered the official barcode of life for the fungal kingdom (6). Unlike the rRNA genes, which have a long history of use as phylogenetic markers, being relatively highly conserved and universally present in all domains of cellular organisms, the non-coding ITS domain has a weak phylogenetic signal due to its hypervariability, making alignment and phylogeny estimation from distantly related clades challenging (7, 8). This limitation impedes the use of phylogenetic diversity metrics for studying fungal community ecology with ITS sequences.

Moreover, phylogenetic reconstruction can be a computationally intensive process, even from short sequence reads (e.g., 16S rRNA gene V4 domain sequences) that yield relatively imprecise phylogenies. This creates a significant bottleneck in terms of time cost and computational resources, which can be limiting, e.g., in resource-limited settings as well as in commercial or clinical settings when turnaround time is an important factor. The primary application of phylogenetic metrics in microbiome research, e.g., relying on short sequence reads to build imprecise phylogenies, is to weight diversity metrics based on the similarity between their constituent sequences, not to infer precise evolutionary relationships per se.

K-mer counting provides an alternative, promising solution for incorporating subsequence information into diversity estimates and microbiome analysis. K-mers consist of the subsequences of length k that compose a given sequence, and k-mer frequencies (as well as sketching/minimization techniques based on k-mer frequencies) have been applied in diverse areas of bioinformatics and microbiome research, e.g., for taxonomic classification (9–12); phylogeny estimation (13); (meta)genome assembly, binning, and comparison (14); and supervised learning (15, 16) in both marker-gene sequence (e.g., 16S rRNA gene) and whole metagenome sequence analysis (17–20). However, k-mer counting has been largely neglected as an approach for feature processing prior to diversity analyses with different alpha and beta diversity metrics, and k-mer counting approaches have yet to be widely adopted in microbiome research.

In this work, k-mer counting was tested as a possible solution for incorporating subsequence-level information into diversity estimates and supervised learning from DNA marker-gene sequence surveys. Results demonstrate that k-mers provide meaningful information about sequence composition, enabling incorporation of this information into diversity metrics without the high computational cost of phylogeny estimation. This suggests that k-mer-based metrics can be used alongside classical metrics as well as phylogenetically aware metrics for quantification of diversity in complex microbial communities.

## Methods

### K-mer counting

K-mer counting is performed from an input series of sequences (in FASTA format) using the CountVectorizer or optionally TfidfVectorizer methods implemented in scikit-learn (21). Matrix multiplication with numpy (22) is then used to multiply the observed frequency of input sequences by the counts of their k-mer constituents to yield a feature table of k-mer frequencies per sample. When the TfidfVectorizer method is used, k-mers are weighted by term frequency-inverse document-frequency (TF-IDF) (23) as a method to upweight k-mer signatures that are more specific to certain classes while downweighting k-mers that are common across classes and hence have little predictive value. The resulting method is implemented in the QIIME 2 (24) plugin q2-kmerizer (https://github.com/bokulich-lab/q2-kmerizer).

### Benchmarking data and data preparation

The Earth Microbiome Project (EMP) (1) dataset was downloaded from the EMP FTP site. The “emp_deblur_150bp.subset_2k” dataset (.biom md5sum = a135e5d53229bf68cb3921fd29b87531) was used, consisting of 16S rRNA gene V4 domain sequences that have been denoised to obtain unique amplicon sequence variants (ASV). To obtain a reasonably sized test dataset, all samples were evenly rarefied to 1000 sequences per sample using the QIIME 2 plugin q2-feature-table (24). This yielded a total of 975 samples that passed these criteria. K-mer frequencies were counted from rarefied feature tables using both CountVectorizer and TfidfVectorizer with the parameters max_df=0.95 and min_df=10 to ignore k-mers that were observed in >95% of ASVs or in fewer than 10 ASVs.

The Global Soil Mycobiome Consortium (GSM) data (2) were downloaded from the PlutoF data repository (https://doi.org/10.15156/BIO/2263453), consisting of full-length fungal internal transcribed spacer (ITS) sequences obtained using PacBio sequencing and clustered into OTUs (however, these are referred to as ASVs in the text for simplicity and consistency). Samples were filtered to remove any with suspected mold or other contamination as indicated in the study metadata. RESCRIPt (25) was used to evaluate sequence length distribution and remove outliers (< 300 nt and > 800 nt long). To obtain a representative subsample, 100 samples were randomly sampled from each of the 7 most abundant biomes represented in the dataset (i.e., those with > 100 samples to exclude undersampled biomes) to yield a test set consisting of 678 samples out of a total of 3,200 samples (678 < 700 because some of the representative samples were subsequently filtered due to the filtering criteria). Rarifying and k-mer counting was performed as described for the EMP dataset.

For plotting prevalence vs. abundance curves, ASV and k-mer abundance was calculated as the total number of observations in each dataset (after rarefying). Prevalence was calculated as the fraction of samples in which that feature was observed.

### Diversity analysis

Alpha diversity (observed features, Shannon entropy, Faith’s Phylogenetic Diversity) and beta diversity (Bray Curtis dissimilarity, Jaccard distance, Aitchison distance, un/weighted UniFrac (3)) were calculated using QIIME 2 version 2024.5 with the q2-diversity plugin. Non-phylogenetic metrics (observed features, Shannon, Bray Curtis, Jaccard, Aitchison) were calculated on rarefied feature tables or k-mer frequency tables. Phylogenetic metrics were calculated only on ASV frequencies, using a phylogeny estimated by fasttree (26) based on a multiple sequence alignment of the ASVs with mafft (27). PERMANOVA tests (28) were performed using the q2-diversity plugin to test for significant differences in beta diversity between EMP ontology level 3 (EMPO 3) categories, which categorize samples based on general sample types. ANOVA tests with false discovery rate (FDR) correction for multiple tests, as implemented in q2-longitudinal (29), were used to test for significant differences in alpha diversity.

To test goodness of fit between PCoA coordinates calculated for each metric and feature processing method (k-mers vs. ASVs), Procrustes analysis (30) was performed using the q2-diversity plugin. Procrustes analysis uses translation, rotation, and uniform scaling to find the best fit between two objects (in this case PCoA coordinates) that minimizes the sum of squares of pointwise differences (the M^2^ statistic reported; lower indicates a better fit). Mantel tests were performed to test the correlation between pairwise distances calculated by each distance metric and feature processing method; Spearman correlation was used to correlate these distances and a higher Rho value indicates a closer correlation. Alpha diversity correlations between metrics and methods were measured using Pearson correlation and Spearman correlation implementations in scipy (31). Alpha correlation and beta diversity principal coordinate analysis (PCoA) plots were generated using the seaborn package (32).

### Supervised learning

Random Forest classifiers (33) with 500 trees were trained using 10-fold nested cross-validation using the QIIME 2 plugin q2-sample-classifier (34). Models were trained on rarefied ASV or K-mer frequency tables and tested on a hold-out set at each fold. Accuracy was measured at each fold by comparing true versus predicted values. Receiver operating characteristic (ROC) curves were calculated for each model using the predicted class probabilities, and used to measure area under the curve (AUC). K-nearest neighbors models (k=3, 10-fold cross-validation) were used to test predictive accuracy for sample classification based on pairwise UniFrac distances. Although this is a different model and approach, it allows comparison of predictive accuracy between ASV or K-mer frequencies versus UniFrac diversity estimates for predicting sample metadata classes.

### Computational performance

Runtimes were measured using a single core on a Macbook Pro with a 2 GHz Quad-Core Intel Core i5 processor and 16 GB RAM. The EMP and GSM datasets were used to provide test data from 150nt long 16S rRNA gene V4 sequences and ∼800nt long PacBio sequences, respectively. Sub-samples of 100, 500, 1000, and 10000 sequences were taken from each dataset to test performance with different sized queries. K-mer counting was performed as described above with TFI-DF and 5000 maximum features. The “align_to_tree_mafft_fasttree” action in the QIIME 2 plugin q2-phylogeny was used to perform pairwise alignment with mafft (27) followed by phylogeny estimation with fasttree2 (26).

### Data and code availability

The EMP and GSM datasets are publicly available as described above. The EMP files used in this study, as well as a subset of the GSM dataset used here (selected as described above), are available on Zenodo (https://zenodo.org/records/13304967). A method for k-mer counting is implemented in the QIIME 2 plugin q2-kmerizer (https://github.com/bokulich-lab/q2-kmerizer).

## Results

The Earth Microbiome Project (EMP) dataset was selected for testing here as a significant global dataset representing most key sample categories, consistently categorized using the EMP ontology, EMPO. This dataset consists of 16S rRNA gene V4 domain amplicon sequence variants (ASVs).

### K-mer diversity corresponds to phylogenetically aware diversity estimates

K-mer counting yields beta diversity ordinations that closely fit to those derived from the phylogenetically aware beta diversity metric UniFrac (Figure 1, Table 1; Procrustes M^2^ = 0.231). This correlation is stronger than the correspondence between k-mer frequency-based ordinations and ASV-based ordinations using non-phylogenetic diversity metrics (M^2^ = 0.365), as well as between ASV-based ordinations and Unifrac-based ordinations, which only show weak correspondence (M^2^ = 0.455). Ordinations based on k-mer frequencies are qualitatively more differentiable between sample classes in the EMP dataset (Figure 1), and yield larger PERMANOVA R^2^ values for differentiating EMPO 3 categories, indicating better quantitative differentiation between sample classes than for ordinations based on ASV frequencies when using the same metrics (Table 2). The k-mer frequency-based PERMANOVA R^2^ values for unweighted (Jaccard) and weighted (Bray Curtis) diversity metrics are also in close correspondence with those for unweighted and weighted UniFrac, respectively (Table 2), indicating that k-mer frequencies and phylogeny-aware metrics have a similar degree of efficacy for differentiating communities based on subsequence-level information, as can be expected from the closely correlated ordinations (Table 1). Bray-Curtis distances based on k-mer frequencies were also closely correlated with weighted UniFrac distances between the same sample pairs (Spearman R = 0.784, P < 0.001) and ordinations based on these distances demonstrated a moderately good fit (Procrustes M^2^ = 0.276).

**Figure 1.**
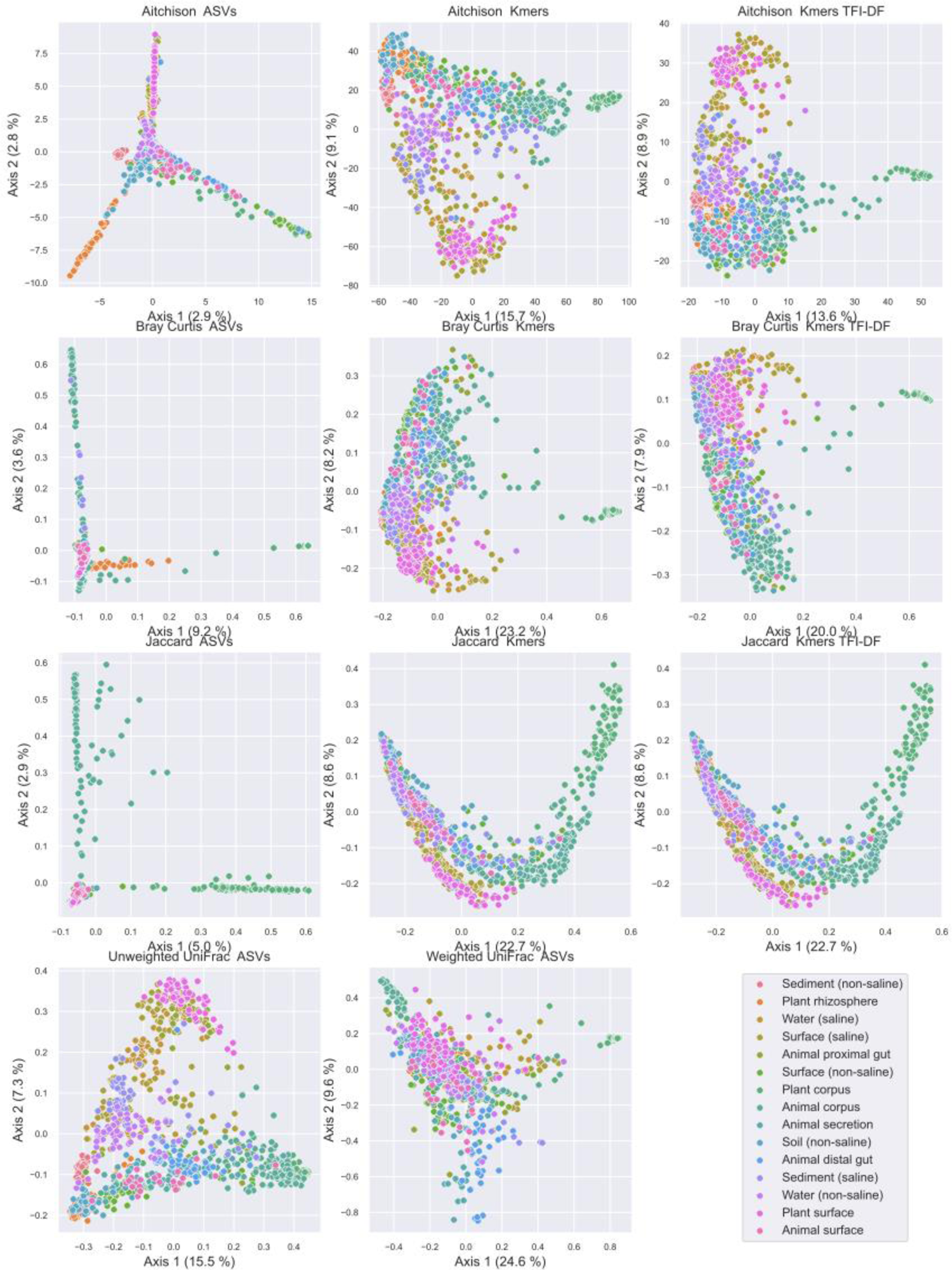
Principal coordinate analysis comparison of sample compositions based on different beta diversity metrics and features (ASVs vs. k-mers). Samples are colored by EMPO 3 category. Axis labels indicate % variance explained. Note that Jaccard distances are identical for k-mer profiles with and without TF-IDF, as TFI-DF was used to weight k-mers based on importance, but not for feature selection. Hence, both profiles contain the same K-mers so binary metrics (e.g., Jaccard) yield identical distances, but weighted metrics (e.g., Bray Curtis) vary due to the TFI-DF weightings.

**Table 1.** Mantel test correlations and Procrustes goodness-of-fit (M^2^) between different distance metrics and feature processing methods.

| | | Mantel<br>Spearman *R* | p-value | Procrustes $M^2$ | p-value |
| --- | --- | --- | --- | --- | --- |
| ASV Jaccard | Unweighted UniFrac | 0.436 | 0.001 | 0.455 | 0.001 |
| ASV Jaccard | K-mer Jaccard | 0.451 | 0.001 | 0.365 | 0.001 |
| K-mer Jaccard | Unweighted UniFrac | 0.396 | 0.001 | 0.231 | 0.001 |
| ASV Bray Curtis | K-mer Bray Curtis | 0.428 | 0.001 | 0.344 | 0.001 |
| ASV Bray Curtis | Weighted UniFrac | 0.360 | 0.001 | 0.519 | 0.001 |
| K-mer Bray Curtis | Weighted UniFrac | 0.784 | 0.001 | 0.276 | 0.001 |

**Table 2.**
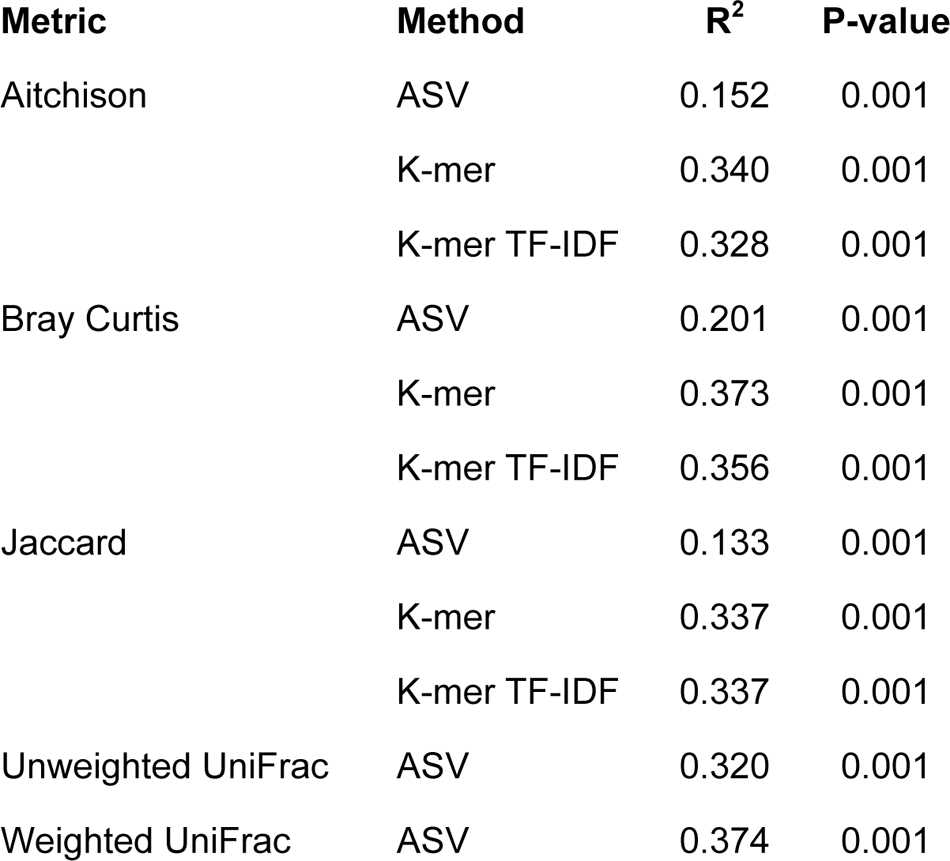
PERMANOVA tests for differentiation by EMPO 3 class.

| Metric | Method | $R^2$ | P-value |
| --- | --- | --- | --- |
| Aitchison | ASV | 0.152 | 0.001 |
|  | K-mer | 0.340 | 0.001 |
|  | K-mer TF-IDF | 0.328 | 0.001 |
| Bray Curtis | ASV | 0.201 | 0.001 |
|  | K-mer | 0.373 | 0.001 |
|  | K-mer TF-IDF | 0.356 | 0.001 |
| Jaccard | ASV | 0.133 | 0.001 |
|  | K-mer | 0.337 | 0.001 |
|  | K-mer TF-IDF | 0.337 | 0.001 |
| Unweighted UniFrac | ASV | 0.320 | 0.001 |
| Weighted UniFrac | ASV | 0.374 | 0.001 |

Supervised learning was also used to determine the predictive value offered by different feature processing methods (ASVs vs. k-mers generated from those ASVs) as well as vs. UniFrac distance estimates from ASVs. Random Forest classification of EMPO 3 categories (using 10-fold nested cross-validation with 500 trees at each fold) demonstrated that predictive accuracy was marginally improved when models were trained on K-mers either with (accuracy = 0.848) or without TFI-DF (accuracy = 0.849) compared to ASVs (accuracy = 0.813), though ROC AUC was equivalent for the three approaches (micro-averaged AUC = 0.99). This exceeded accuracy for k-nearest neighbors classification of EMPO 3 categories based on weighted UniFrac (accuracy = 0.782) and unweighted UniFrac (accuracy = 0.825). Hence, the expanded feature space offered by k-mer frequencies may increase classification accuracy in some contexts; further optimization (e.g., through feature selection) could be done to further improve performance, but will be dataset dependent and is out of scope in this work. In any given experiment, ASVs and k-mer frequencies both are recommended as inputs for supervised learning models to determine the predictive power offered in the context of that experiment.

### K-mer length influences diversity estimates

The impact of k-mer length (i.e., n-gram size) on beta diversity ordinations (Figure 2) and Procrustes goodness of fit (Table 3) with phylogenetically aware beta diversity metrics was tested on the same dataset, comparing weighted (k-mer Bray Curtis vs. ASV weighted UniFrac) and unweighted (k-mer Jaccard vs. ASV unweighted UniFrac). For this test, TF-IDF vectorization while constraining the feature table to the top 5000 most important k-mers so that k-mer count is kept constant across all tests. Results indicate that k-mer lengths between k=7 and k=32 yielded similar Jaccard distance ordinations (Figure 2) to UniFrac distance ordinations (Figure 1), with very good fits (Table 3). K-mer lengths of k=7 and k=9 yielded the best fits for weighted metrics (Bray Curtis and weighted UniFrac) (Table 3) and hence k=7 was kept for all other tests in this work. Nevertheless, the influence of k-mer length on diversity estimates may vary by target domain, and depend on the underlying diversity, and hence in any given experimental scenario it is recommended to repeat this process when applying this method in different experimental contexts. However, very small k-mer lengths reduce the feature space overall, leading to a small number of highly prevalent features, resulting in poorer diversity estimates and limited ability to differentiate samples; very large k-mers may be expected to have the same effect.

**Figure 2.**
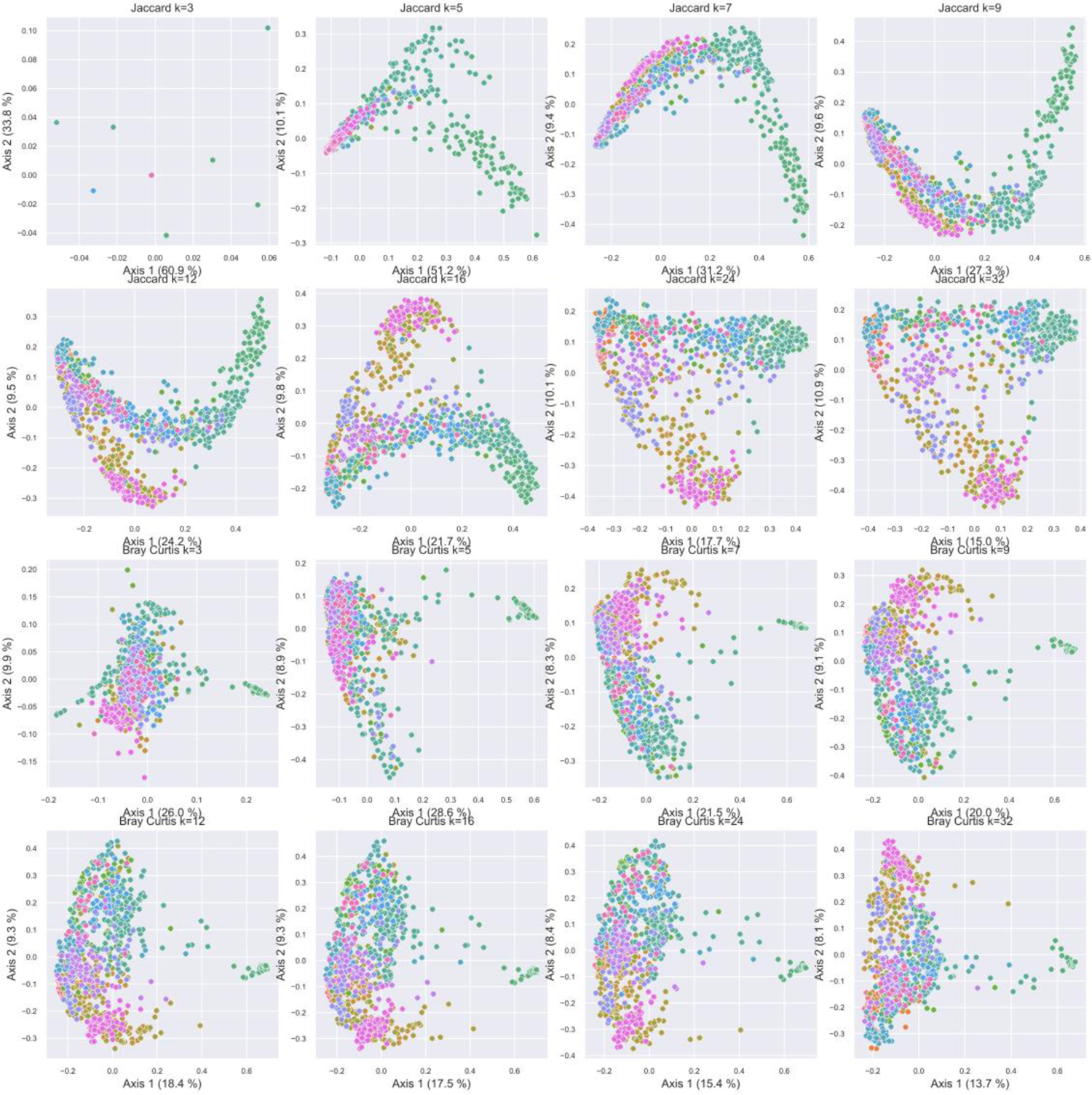
Influence of k-mer length on PCoA ordinations based on Jaccard distance (top two rows) and Bray Curtis dissimilarity (bottom two rows). Samples are colored by EMPO 3 category (see legend in Figure 1). Axis labels indicate % variance explained.

**Table 3.** Procrustes goodness-of-fit (M2) scores by k-mer length (*k*) and metric between k-mer-based distances and UniFrac distances.

| Method* |  |  |
| --- | --- | --- |
| *k* | Unweighted** | Weighted*** |
| **3** | 0.933 | 0.469 |
| **5** | 0.536 | 0.319 |
| **7** | 0.268 | 0.288 |
| **9** | 0.249 | 0.288 |
| **12** | 0.237 | 0.297 |
| **16** | 0.229 | 0.3 |
| **24** | 0.228 | 0.324 |
| **32** | 0.236 | 0.394 |

\*For all tests $P$ values = 0.001
\*\*Unweighted = Jaccard distance on k-mers vs. unweighted UniFrac distance on ASVs.
\*\*\*Weighted = Bray Curtis distance on k-mers vs. weighted UniFrac distance on ASVs.

### K-mer diversity correlates with ASV-based alpha diversity estimates

Next, alpha diversity estimates for the EMP data were compared between 7-mers (observed k-mers, Shannon entropy) and ASVs (observed ASVs, Shannon entropy, and the phylogenetically aware metrics Faith’s phylogenetic diversity). Overall, good correlations were seen between all metrics, indicating that richness, entropy, and phylogenetic diversity are correlated within microbial communities on a global scale (i.e., within the EMP dataset), and that k-mer diversity is closely correlated to ASV diversity, though the relationship is slightly non-linear (Figure 3, Table 4). In addition, the closest correlation observed between any of the two metrics tested was between observed k-mers and Faith’s phylogenetic diversity (Spearman *R* = 0.997, *P* < 0.001; Table 4), indicating that these metrics have a very close (and slightly non-linear) correlation.

**Figure 3.**
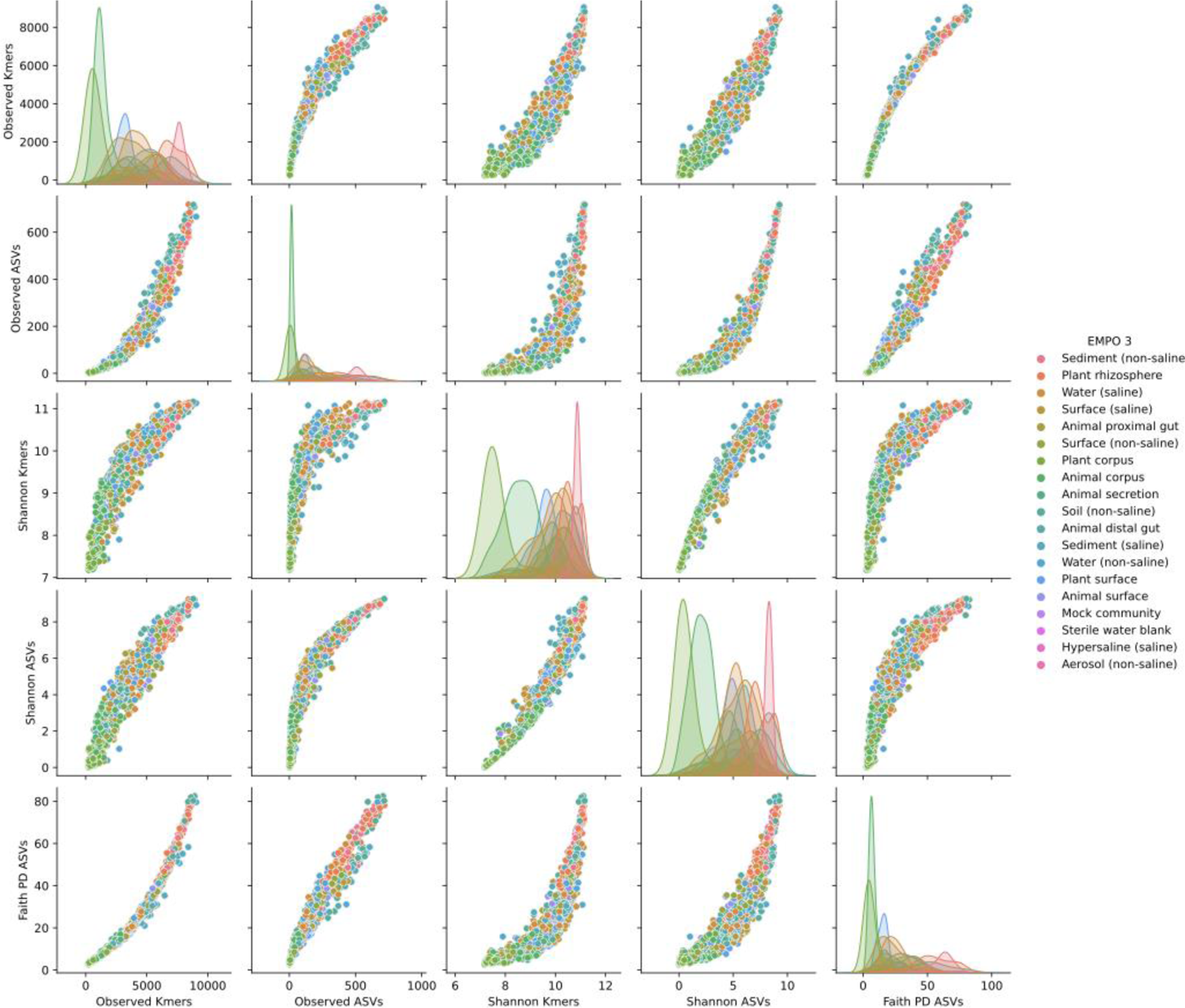
Pairplot of correlations between k-mer- and ASV-based alpha diversity estimates.

**Table 4.** Correlations between alpha diversity estimates.

|  |  | Pearson *R* | P-value | Spearman *R* | P-value |
| --- | --- | --- | --- | --- | --- |
| Observed ASVs | Observed k-mers | 0.941 | < 0.001 | 0.987 | < 0.001 |
| Shannon ASVs | Shannon k-mers | 0.971 | < 0.001 | 0.972 | < 0.001 |
| Observed ASVs | Faith PD (ASVs) | 0.975 | < 0.001 | 0.983 | < 0.001 |
| Faith PD (ASVs) | Observed k-mers | 0.968 | < 0.001 | 0.997 | < 0.001 |
| Shannon ASVs | Faith PD (ASVs) | 0.888 | < 0.001 | 0.953 | < 0.001 |
| Faith PD (ASVs) | Shannon k-mers | 0.828 | < 0.001 | 0.948 | < 0.001 |

Hence, k-mer diversity could be used in addition or instead of phylogeny-based diversity metrics as a measurement of subsequence diversity present within a community. This comes with the added advantage that k-mer-based metrics can be directly compared between datasets when parameters (e.g., rarefaction depth and k-mer length) are kept consistent, whereas phylogenetically aware metrics are only comparable when an identical phylogeny was used, which can limit comparability of results in the literature. However, this comes at the cost of interpretability; whereas richness measurements are easily interpretable (e.g., number of unique species or sequence variants observed in a sample), k-mer richness does not map to such an intuitive value, though arguably this is still more intuitive and interpretable than for phylogenetic alpha diversity, e.g., cumulative amount of branch length on a tree (depending on the phylogeny and the unit of measurement).

### K-mer diversity incorporates subsequence information into fungal ITS diversity estimates

To evaluate the utility of k-mer counting in fungal ITS studies, the Global Soil Mycobiome (GSM) dataset was selected as a test dataset, consisting of PacBio full-length fungal ITS sequences from soil microbiomes collected from different locations, categorized according to biome.

Different k-mer lengths were tested to determine how this impacted weighted (Bray Curtis dissimilarity) and unweighted (Jaccard distance) beta diversity in the GSM dataset. Low k-mer lengths led to unweighted distances that were distorted with outliers and weighted metrics appeared less sensitive to k-mer length (Figure 4). K-mer lengths of k=7 and k=9 still led to highly distorted Jaccard distances, suggesting that rare k-mers found only in some sequences and samples could cause this distortion. K-mer prevalence-abundance curves (Figure 5) indicate that smaller k-mer lengths lead to fewer highly prevalent features, leading to poor discriminatory power. Larger k-mer lengths led to a larger number of k-mers with more variable prevalence, leading to more favorable ordinations and discriminatory power. Thus, a k-mer length of 16 was chosen for further analyses with the GSM dataset.

**Figure 4.**
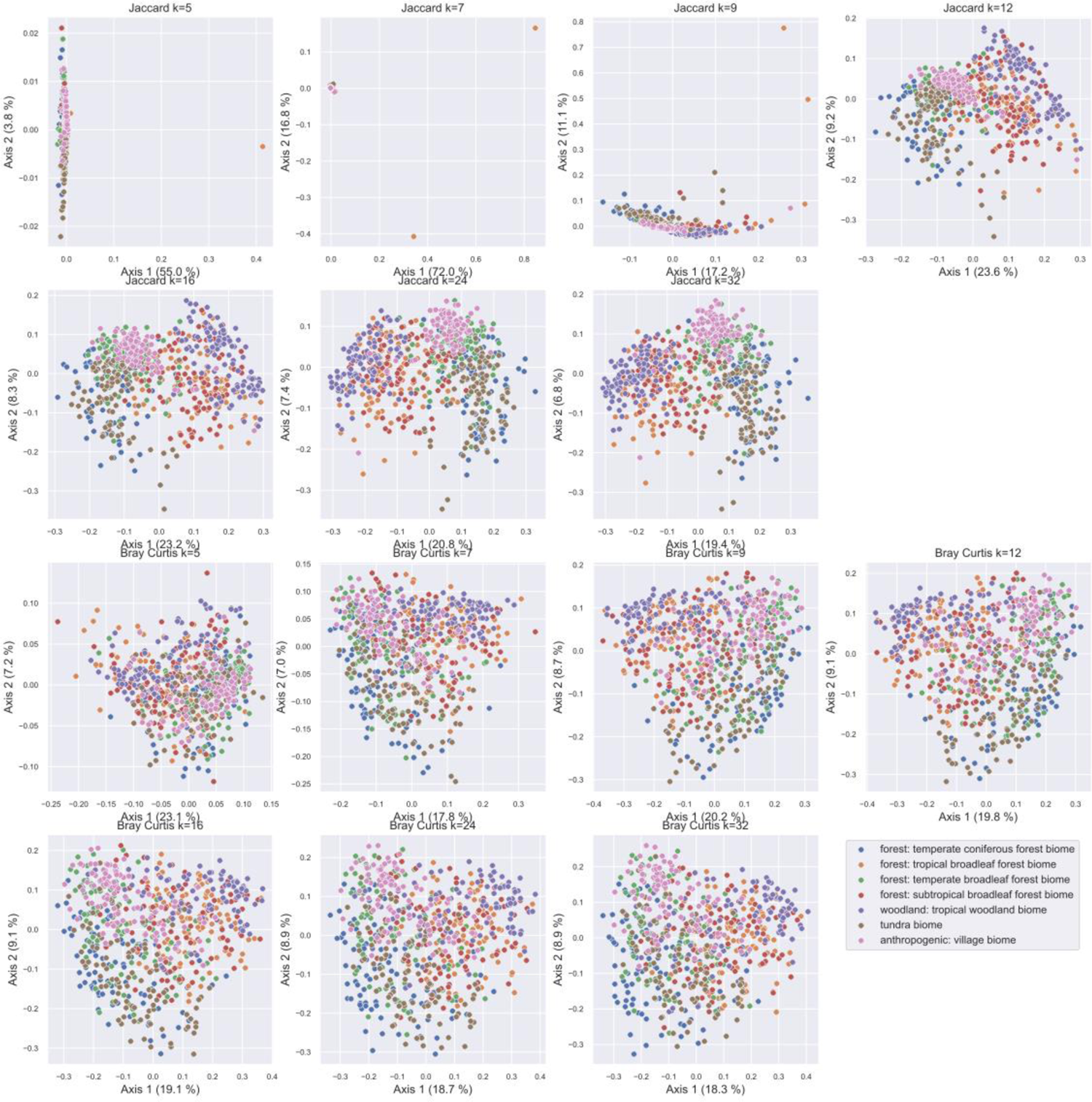
Influence of k-mer length on PCoA ordinations based on Jaccard distance (top two rows) and Bray Curtis dissimilarity (bottom two rows) in the Global Soil Mycobiome dataset. Samples are colored by biome category. Axis labels indicate % variance explained.

**Figure 5.**
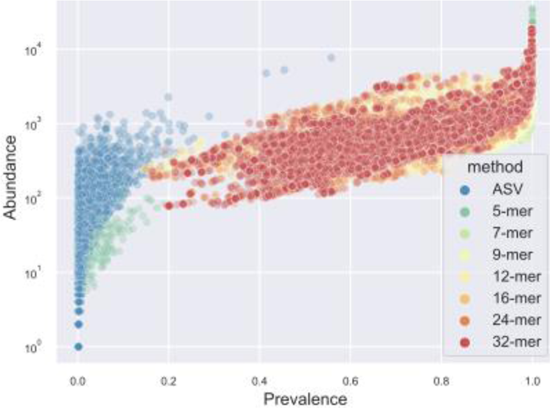
Scatter plot of feature prevalence vs. abundance for ASVs and k-mers of different length in the Global Soil Mycobiome dataset.

Beta diversity was calculated as described for the EMP dataset, comparing ASV-based vs. K-mer-based diversity using weighted (Bray Curtis dissimilarity), unweighted (Jaccard distance), and compositionally aware metrics (Aitchison distance) (Figure 6). The ASV-based PCoA ordinations all demonstrated good separation of most sample types, but with very low percentage variance explained (maximum 5.7% on the first two PCs) and highly distorted plots with poor separation of several sample classes (i.e., biome types). On the other hand, k-mer-based PCoA ordinations all demonstrated good separation of sample types, high percentage variance explained (between 25.7% to 30.7% in the first two PCs), and smooth, undistorted ordinations. Hence, k-mer counting, when correctly parameterized, appears to be an effective strategy to incorporate subsequence-level information into community diversity estimates based on ITS sequences, leading to better discriminatory power than ASV (or OTU-based) methods, which do not incorporate sequence similarity into their estimates, and hence treat all features as equally unrelated entities. This limits discriminatory power, or may even artificially amplify differences between samples, e.g., when differences between samples are driven by genetically similar organisms (e.g., different strains within the same species). For this reason, phylogenetically aware metrics have been recommended for use alongside classical beta diversity metrics to evaluate how genetic diversity influences beta diversity (3). In lieu of a robust method for measuring phylogenetic diversity with ITS sequences (as a non-coding, phylogenetically uninformative region), k-mer counting provides a solution for incorporating subsequence-level information into diversity estimates to fill this niche. This is not intended as a replacement for ASV- or OTU-based diversity estimates, but as a complimentary approach.

**Figure 6.**
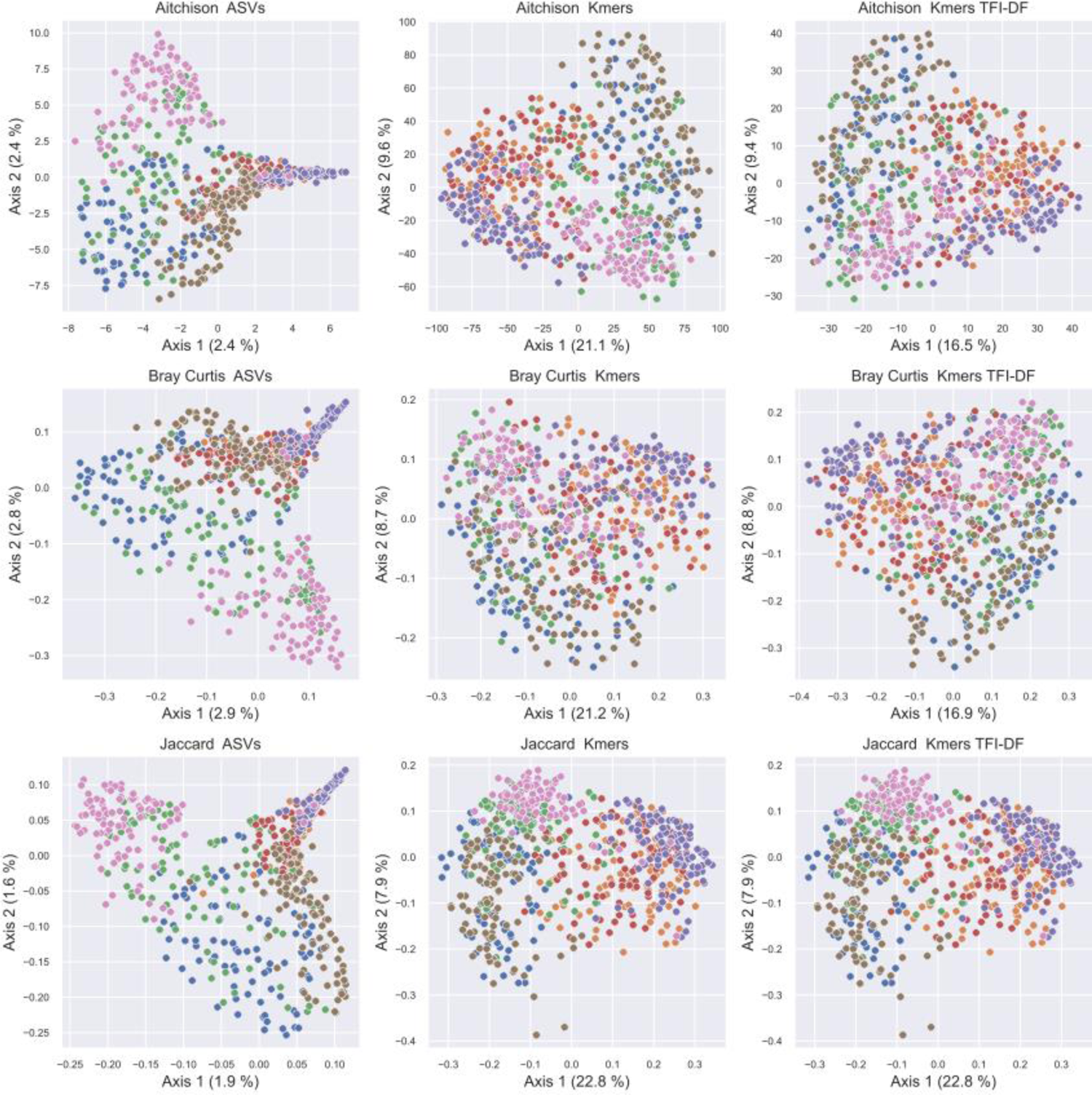
Principal coordinate analysis comparison of sample compositions based on different beta diversity metrics and features (OTUs vs. k-mers) in the Global Soil Mycobiome dataset. Samples are colored by biome category (See Figure 4 for legend). Axis labels indicate % variance explained. Note that Jaccard distances are identical for k-mer profiles with and without TF-IDF, as TFI-DF was used to weight k-mers based on importance, but not for feature selection. Hence, both profiles contain the same K-mers so binary metrics (e.g., Jaccard) yield identical distances, but weighted metrics (e.g., Bray Curtis) vary due to the TFI-DF weightings.

Next, alpha diversity was measured using several metrics based on both ASVs and k-mer frequency. As ITS is a non-coding, phylogenetically uninformative region, it would not be meaningful to calculate phylogenetically aware diversity metrics (e.g., UniFrac or Faith’s PD) with the GSM fungal ITS dataset for comparison vs. k-mer-based diversity estimates as shown above for the EMP dataset to demonstrate close correspondence between these methods. Thus, with the GSM dataset the next analysis was instead to examine how ASV- or k-mer-based alpha diversity estimates lead to different conclusions about community-level diversity. Results demonstrate that k-mer-based diversity estimates have lower relative variance than ASV-based diversity estimates (Figure 7), leading to better discrimination between groups and more significant pairwise differences (Figure 8).

**Figure 7.**
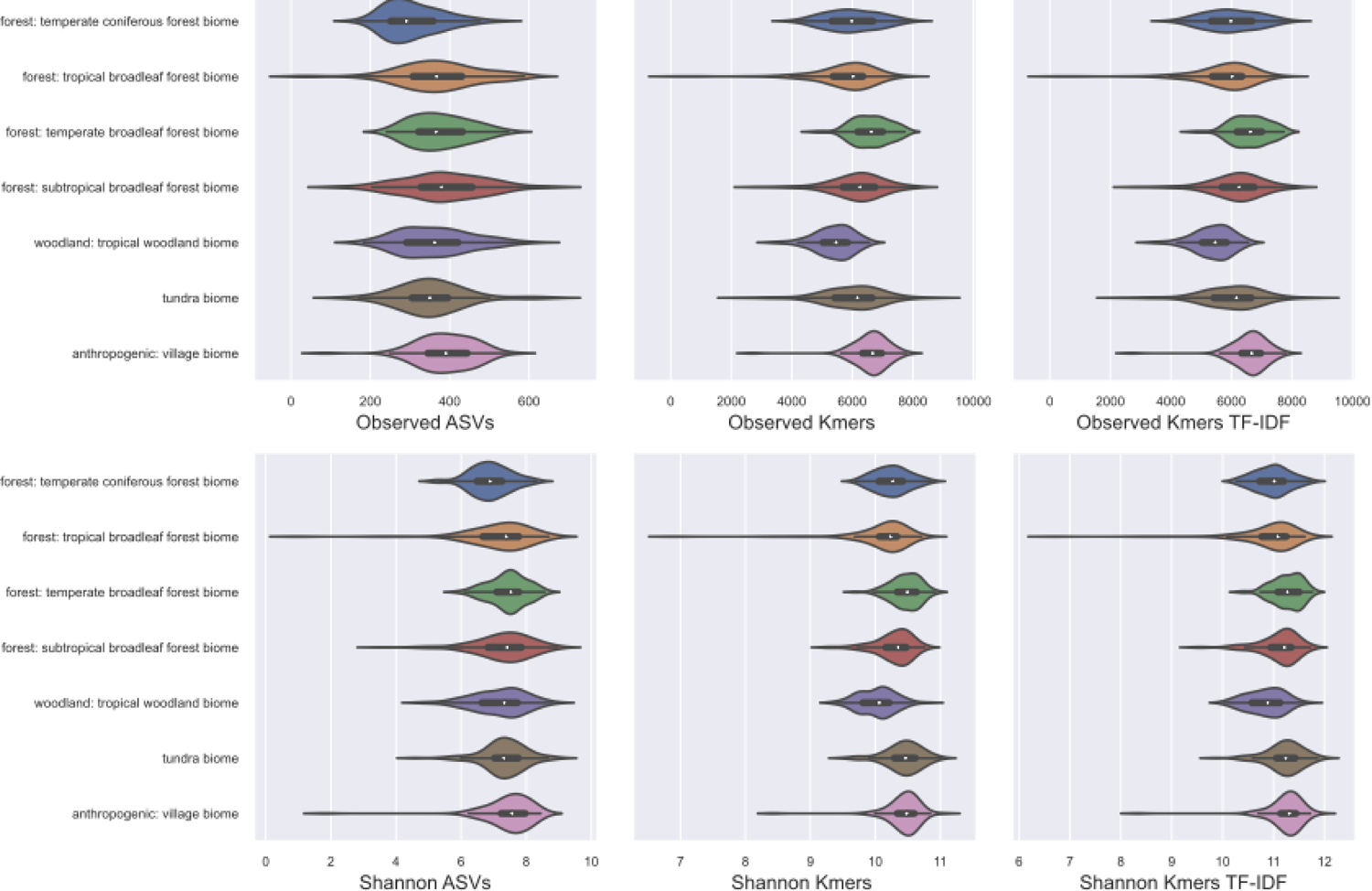
Alpha diversity estimates for the top seven biomes in the Global Soil Mycobiome dataset.

**Figure 8.**
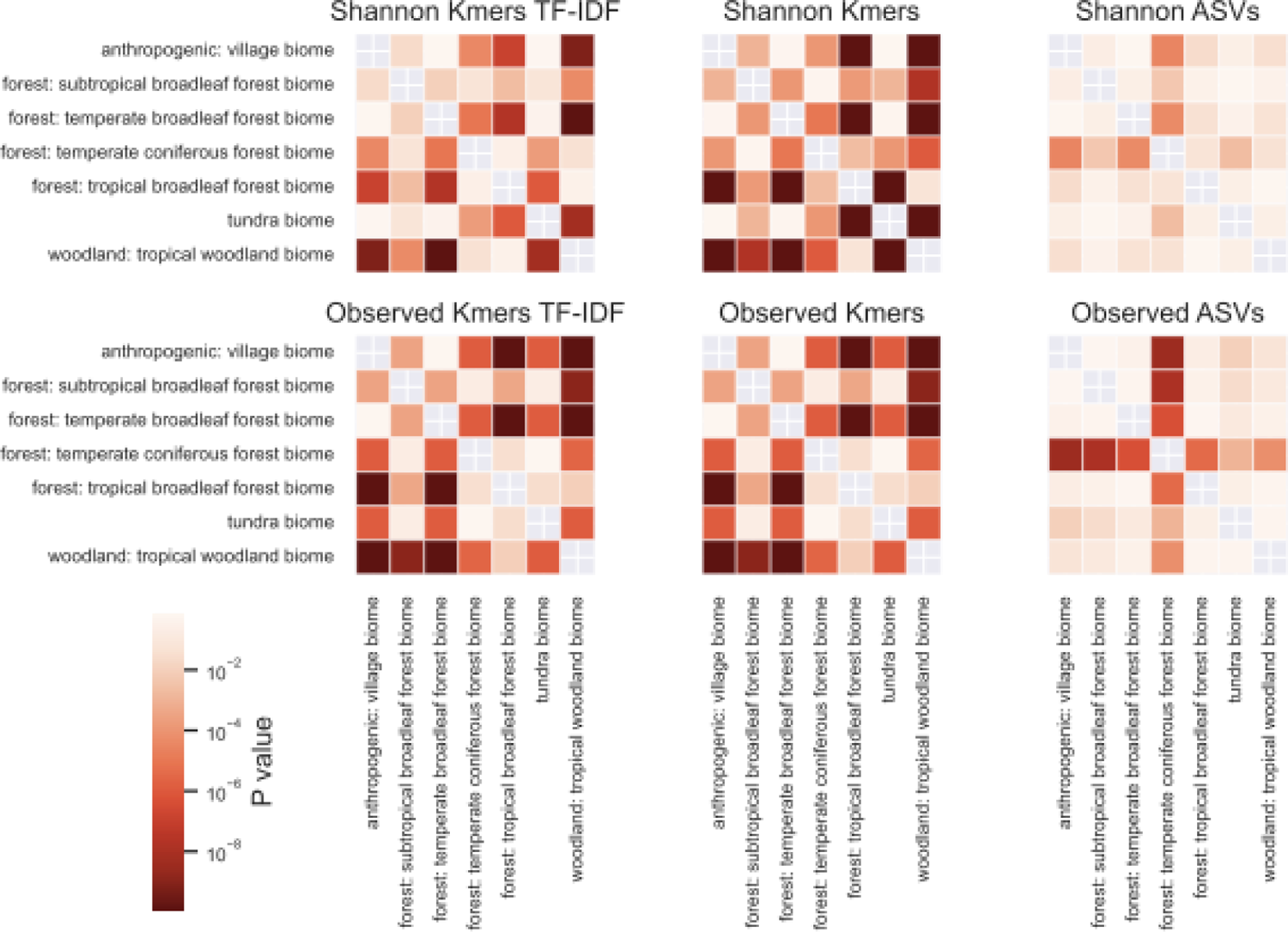
K-mer-based alpha diversity metrics lead to more significant differences between biomes in the Global Soil Mycobiome dataset. Heatmaps depict the False-discovery-rate-corrected P-values for pairwise ANOVA tests between each biome.

This test does not imply that k-mer-based metrics are superior to classical ASV-based metrics, or should serve as a replacement. Instead, these should be used as complementary approaches, similarly to how phylogenetic alpha diversity metrics are used as complementary metrics for 16S rRNA gene diversity. The underlying features are fundamentally different and would not be expected to yield equivalent results, but instead give complementary views of the diversity. Importantly, ASV richness (observed ASVs) has the advantage of being easily interpretable. Hence, these are proposed as complementary, not competing approaches for feature processing. Moreover, k-mer counting fills the niche for incorporating subsequence-level information into fungal ITS diversity metrics (both alpha and beta), and here demonstrates how such information can lead to different interpretations vs. ASV (or OTU-based) metrics.

### K-mer counting is computationally inexpensive

One motivation for k-mer counting is to provide a computationally inexpensive process for incorporating sequence information into diversity estimates. Precise phylogeny estimation (and the pairwise sequence alignment step that precedes it) can be very computationally intensive on large datasets, creating a significant resource and time bottleneck for many researchers. Runtimes were compared between k-mer counting and pairwise alignment with mafft (27) followed by phylogeny estimation with fasttree2 (26). Runtimes were calculated as a function of sequence count (100, 500, 1000, or 10000 sequences) and sequence length (150nt long 16S rRNA gene V4 sequences from EMP or ∼800nt long PacBio sequences from GSM).

Results show that k-mer counting is computationally inexpensive, scaling linearly and taking seconds to process 10000 sequences and their frequencies in hundreds of samples from the EMP and GSM datasets, compared to many minutes to over an hour for alignment and phylogeny estimation with mafft and fasttree2 (Figure 9). As expected, the longer GSM sequences led to significant increases in runtime for mafft + fasttree, but only a minimal increase for k-mer counting. This indicates that k-mer counting takes between 1 and 3 orders of magnitude less runtime on equivalent resources, compared to mafft + fasttree, depending on the number and length of sequences. Moreover, this is just a 10k sequence subset of the EMP and GSM datasets; the full studies reported 307,572 and 722,682 unique sequences, respectively, which would lead to significantly longer runtimes for datasets of that scale. However, all tests were performed with a single core for the purposes of benchmarking, and so in practice all runtimes could be reduced via parallelization when sufficient resources are available.

**Figure 9.**
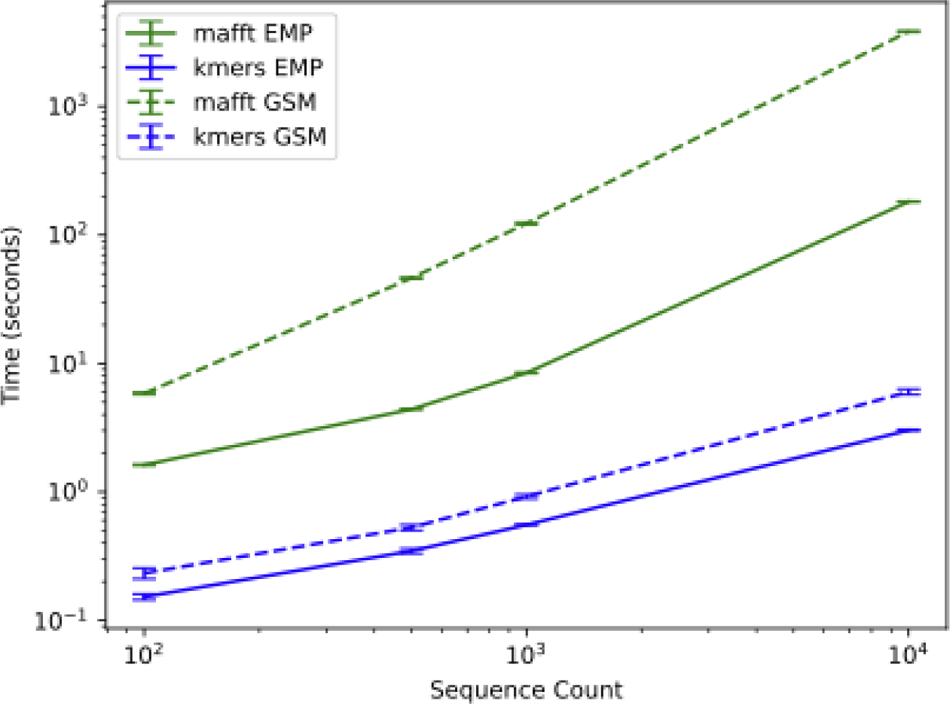
Analysis runtime (seconds) for mafft pairwise alignment + fasttree2 phylogeny estimation (green) versus k-mer counting (blue) as a function of sequence count and length. The X and Y axes are displayed in logarithmic scale.

## Discussion

K-mer frequency counting is proposed here as an approach for sequence processing prior to alpha diversity, beta diversity, and supervised learning analyses in sequence-based microbiome profiling. This is proposed as a complementary method, not as a replacement for classical approaches for diversity measurements. K-mer counting offers several advantages:

1. K-mer counting allows rapid diversity comparison between samples, yielding ordinations that are closely correlated to those derived from the phylogenetically aware beta diversity metric UniFrac (Figure 1; M^2^ = 0.231, P < 0.001). Conventional non-phylogenetic beta diversity estimates are calculated based on species or sequence variant counts, which effectively treats all features as being fully independent and equidistant from one another. This has been an advantage of phylogenetically aware methods, which compares samples based on the amount of genetic diversity present in each sample, e.g., the fraction of branch length in a phylogeny that two samples have in common (3). K-mer counting offers this same advantage, comparing samples based on subsequence-level diversity (i.e., shared or dissimilar k-mers) while avoiding the high computational cost and challenges of accurate phylogeny estimation. This corresponds to the findings of Zhai and Fukuyama (20), who also found that Euclidean distances based on k-mer spectra of whole metagenome data correlated closely to the phylogenetic metric Rao’s quadratic entropy. Together, this suggests that k-mer-based distance metrics are a powerful tool for diversity measurements from both marker gene and metagenome datasets, which incorporate pseudo-phylogenetic information into diversity measurements as well as supervised learning analyses.
2. Runtime is significantly less than phylogeny estimation techniques (Figure 9). Nevertheless, k-mer counting is not intended as a phylogenetic estimate or replacement of phylogeny estimation wherever accurate phylogenies are required. However, in the context of microbial community beta diversity metrics, the goal is frequently to differentiate samples based on overall genetic diversity, not to estimate accurate phylogenies per se. Hence, k-mer counting is useful for distinguishing samples based on sequence topology.
3. This approach allows sequence-based alpha and beta diversity comparison between non-coding regions with high mutation rates and poor phylogenetic signal, e.g., ITS sequences, which are otherwise challenging to use with phylogenetically aware beta diversity techniques.
4. This approach can be used with any standard diversity metric, providing a high degree of flexibility and compatibility with different experimental hypotheses. Hence, k-mer frequencies can be compared with both qualitative (e.g., Bray-Curtis dissimilarity) and quantitative metrics (e.g., Jaccard distance). Moreover, in contrast with existing phylogenetically aware methods, it is compatible with compositionally aware beta diversity metrics (e.g., Aitchison distance).
5. The method is totally reference-free, enabling use in untargeted analysis across a range of ecosystems and target sequences. In other words, no reference data are used when counting k-mers, eliminating the need for curated reference sequences, alignments, or phylogenies in this type of analysis. This contrasts with some phylogeny estimation approaches, including insertion-based methods (5), which often rely on a reference alignment or phylogeny for reliable placement, limiting use for some sequence domains (e.g., ITS sequences and many non-ribosomal domains).

However, k-mer counting has several limitations that should be considered when applying this method:

1. This approach should be used to compare sequences from the same target region. Comparing k-mer frequencies from different marker genes or domains may not yield meaningful results, as the k-mer frequency is expected to differ between different genes and domains. On the other hand, k-mer counting has been demonstrated as an effective technique for comparing whole (meta)genomes (17–20), so remains scalable as long as comparable methodology was used, and potentially would provide a means of integrating datasets (an application out of scope in the current work and will be explored in the future).
2. K-mer counting as implemented here is not a replacement for phylogenetic techniques, i.e., when an accurate estimate of evolutionary relationships is required, though k-mer counting has been demonstrated to accurately reflect evolutionary distances between genome sequences (13). Further work would be needed to demonstrate such an application with short DNA marker genes.
3. K-mer counting can significantly expand the feature space, as each original sequence is broken up into many constituent k-mers. For this reason, various parameters are exposed to the user to constrain the feature space by filtering k-mers based on the minimum or maximum number of observations; specifying a maximum number of k-mers; and using TF-IDF to rank k-mers by importance. As many k-mers will also be highly redundant between different sequence variants (especially for relatively highly conserved sequence domains), these parameters are recommended to reduce redundancy as well as computational demands when appropriate.

In conclusion, k-mer counting appears to be a suitable and efficient strategy for feature processing prior to diversity estimation as well as supervised learning in microbiome surveys. This allows incorporation of subsequence-level information into diversity estimation without the computational cost of pairwise sequence alignment, and is theoretically compatible with any classical (non-phylogenetic) diversity metric. K-mer-based diversity metrics demonstrated close correspondence with weighted and unweighted UniFrac metrics and Faith’s PD across sample types in the global-scale EMP and GSM datasets, suggesting that k-mer frequencies yield approximately similar information to these phylogenetic metrics. K-mer counting is proposed as a complementary approach for feature processing prior to diversity estimation and supervised learning analyses, enabling large-scale reference-free profiling of microbiomes in biogeography, ecology, and biomedical data.

